# Prototyping of *Bacillus megaterium* genetic elements through automated cell-free characterization and Bayesian modelling

**DOI:** 10.1101/071100

**Authors:** Simon J. Moore, James T. MacDonald, Sarah Weinecke, Nicolas Kylilis, Karen M. Polizzi, Rebekka Biedendieck, Paul S. Freemont

## Abstract

Automation and factorial experimental design together with cell-free *in vitro* transcription-translation systems offers a new route to the precise characterization of regulatory components. This now presents a new opportunity to illuminate the genetic circuitry from arcane microbial chassis, which are difficult to assess *in vivo*. One such host, *Bacillus megaterium*, is a giant microbe with industrial potential as a producer of recombinant proteins at gram per litre scale. Herein, we establish a *B. megaterium* cell-free platform and characterize a refactored xylose-repressor circuit using acoustic liquid handling robotics to simultaneously monitor 324 reactions *in vitro*. To accurately describe the system, we have applied a Bayesian statistical approach to infer model parameters by simultaneously using information from multiple experimental conditions. These developments now open up a new approach for the rapid and accurate characterization of genetic circuitry using cell-free reactions from unusual microbial cell chasses for bespoke applications.

## Introduction

Cell-free *in vitro* transcription and translation (ITT) systems utilise cell extracts (1–4) or purified components (5) to synthesise proteins encoded by DNA circuits. Recently, a renaissance in cell-free has captured the imagination of synthetic biologists to explore genetic pathway design (6–9), develop on-site viral biosensors (10), produce antibodies in bioreactors (11) and engineer microfluidic biochip devices (12).

One unexplored area of cell-free protein synthesis is the ability to study the genetic regulatory components of non-model microbial hosts directly in a test tube. Arcane microbial hosts have the potential to provide significant advantages over traditional model organisms and the ability to genetically engineer these microbes is likely to become increasingly important (13). In particular, organisms adapted for growth in extreme environments or able to produce valuable niche biomolecules at high yields from inexpensive substrates are of particular interest (14–17). However, these microbial systems are often not very amenable to genetic manipulation and there is a need to expand the available toolbox of characterized genetic parts. Here, we have developed a cell-extract based ITT system for one such arcane microorganism, *Bacillus megaterium*. We have combined this new ITT system with factorial experimental design, high-throughput automation via acoustic liquid handling robotics and Bayesian model parameter inference methods to create a platform to enable the rigorous and rapid characterization of novel genetic parts. A significant advantage of cell-free systems is that the defined reaction conditions permit the accurate characterization of genetic parts using sophisticated modelling techniques (7, 8, 18, 19). However, such approaches have previously generally been applied to *Escherichia coli* gene expression circuits.

*Bacillus megaterium* is a giant microbe with the potential for biotechnology applications (20, 21). Industrially it has been used to produce penicillin G amidase (22), (β-amylases and vitamin B_12_ (23–25). In comparison to its closely related cousin *Bacillus subtillis*, *B. megaterium* is relatively uncharacterized but provides major advantages such as stable plasmid maintenance (26), minimal neutral-alkaline protease activity (27) and the ability to metabolise low cost precursors (20, 22, 28). Moreover, *B. megaterium* also offers a strong native Sec-dependent secretion apparatus (29, 30), which is a desirable feature for downstream processing, whilst for recombinant gene expression the xylose-inducible promoter system (31–33) produces recombinant proteins to the gram per litre scale (28,30). In addition, protein purification and secretion tools are available (33,34). From these tools, *B. megaterium* has recently been utilised to access challenging proteins (Fe-S, corrinoid enzymes), such as the reductive dehalogenase that were previously insoluble or inactive from using the routine ‘workhorse’ *E. coli*(25, 35–37).

One barrier to the widespread use of *B. megaterium* is a low-efficiency protoplast transformation procedure (38). Our automated ITT and modelling platform enables the rapidly validation of gene expression tools. We tested a designed xylose-repressor circuit by combinatorially varying three experimental conditions (DNA, repressor and inducer concentrations), with the automated generation of transfer instructions for an acoustic liquid handling robot to rapidly set up 324 separate *in vitro* reactions. To our knowledge, this demonstrates the first fully automated platform for μL scale cell-free reaction screens. To model the obtained experimental data, we used a Bayesian statistical inference scheme to rigorously infer unknown ITT kinetic parameters that describe the cell free reaction. The development of a *B. megaterium* ITT system serves as a model for rapidly prototyping novel genetic parts from a relatively uncharacterized host thus providing a basis for future engineering of this model organism. Our platform also provides an integrated high-throughput experimental and modelling framework for characterizing cell extracts from other unusual chassis.

## Methods

### Strains and Plasmids

*B. megaterium* wild-type strain DSM319 was used for preparing the cell-free extract (66), *E. coli* strain BL21 (DE3) Star^TM^-pLysS (LifeTechnologies, Darmstadt, Germany) for the recombinant production of the His_6_-tagged *B. megaterium* DSM319 repressor protein XylR (see Supplementary text) and *E. coli* strain DH10B (Invitrogen, San Diego, USA) for routine cloning. All the plasmids and oligos used in this study are listed in Supplementary Tables S1-3, respectively. For construction of plasmids, please see Supplementary text.

### Growth and recombinant protein production in *B. megaterium*

We utilised an improved fluorescence variant (GFP^+^ - F64L/ S65T/ Q80R/ F99S/ M153T/ V163A) of the wild-type GFP (67), herein simply referred to as GFP, to monitor protein production in *B. megaterium* ITT, along with mCherry. Protoplasted *B. megaterium* cells were individually transformed with the corresponding plasmids and cultivated as described before (68).

### B. megaterium ITT extract preparation and reaction conditions

A *B. megaterium* DSM319 ITT system was developed based on the *E. coli* method of Sun *et al* (40). Full details and modifications are outlined in the Supplementary text and Table S4 This method yielded between 27-34 mg mL^−1^ of cell-free protein extract. All ITT experiments were repeated a minimum of two times, whilst individual assays were collected a triplicate of technical repeats.

### High-throughout automated robotics and experimental set-up

For automation, an Echo®525 liquid handler (Labcyte, Inc.) was combined with a factorial experimental design approach using a script to automate transfer volume instructions and well positions (see Supplementary text). Liquid droplets were transferred as multiples of 25 nL to a final volume of 10 μL as technical triplicate repeat. Plates were sealed with Breathe-Easy^®^ sealing membrane (Sigma) and briefly centrifuged at 1,000 × *g* for 10 seconds. A CLARIOStar plate reader (BMG Labtech, Germany) was used for cell-free incubations and fluorescence measurements. Standard measurements were recorded every 10 min for 40 cycles at 30°C with 10 sec of 250 rpm orbital shaking prior to measurement. Depending on sample size, RNA measurements were recorded 30, 60 or 120 seconds. Fluorescence settings are provided in the SI text.

### Bayesian statistical modelling

The cell-free transcription and translation reactions were modelled using a system of ordinary differential equations (ODEs).

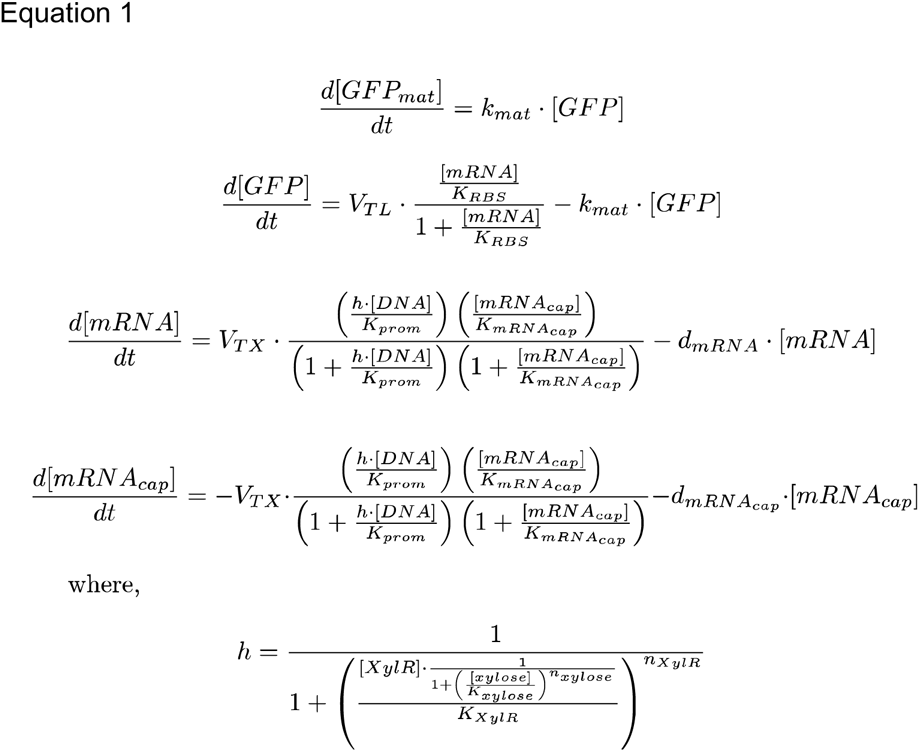

The model consisted of 5 chemical reactions, 4 species (Supplementary Table S5), and 15 parameters (Supplementary Table S6). The regeneration of ATP, GTP, CTP and UTP from the secondary energy source was not explicitly modelled and these were all lumped into a single species, mRNA^cap^ (mRNA capacity). It was assumed that NTPs were also consumed by non-productive side-reactions and this was modelled as a first-order degradation reaction of mRNA^cap^. The transcription rate was modelled assuming multi-substrate Michaelis-Menten kinetics with random order binding of DNA template and mRNA^cap^. For the xylose inducible promoter, the fraction of free DNA was approximated using the Hill equation. The mRNA degradation reaction was assumed to be a first order reaction. Translation rate was modelled using Michaelis-Menten kinetics with mRNA as the only substrate. Amino acids were assumed to be in excess so were not explicitly considered. As in previous models of cell-free reactions, consumption of ATP and GTP during translation was not considered (18). The translation reaction produced non-fluorescent immature GFP and this was assumed to mature in a first-order reaction into the detectable fluorescent species, GFP_mat_. The full system of ODEs is shown in the equation 1.

Parameters and initial values (for [mRNA_cap_]) were inferred using Markov chain Monte Carlo (MCMC) implemented in a C++ program using Boost.odeint library to numerically integrate the system of ODEs. A multivariate normal distribution was used as the proposal distribution during MCMC and moves were accepted following the Metropolis criterion. A uniform or a multivariate Gaussian prior distribution was used for all parameters. Experimental errors were assumed to be independent and normally distributed. The log-likelihood function was therefore defined as the sum of logs of the normal probability density function where the means and standard deviations were taken from the experimentally derived values for each data point. Model parameters were inferred simultaneously from multiple experimental datasets by generating multiple trajectories from the same system of ODEs and evaluating the log-likelihood function. Initial values for all species other than [mRNA_cap_] were set 0. Although initial NTP concentrations in the reaction are known, the overall NTP capacity of the system was unknown as the efficiency of NTP regeneration from the secondary fuel source was unknown. For this reason, the initial value of [mRNA_cap_] was inferred. In most experiments, only the mature GFP concentration was measured. Where the mRNA concentration was experimentally determined, this was also included in the log-likelihood function. Source code is available from https://github.com/jmacdona/B_megaterium_modelling.

## Results

### Isolation of a strong constitutive promoter for *B. megaterium* ITT

To achieve quantitative fluorescence measurements in ITT reactions, a strong promoter system is required. To initially prototype a *B. megaterium* ITT system without regulatory components (e.g. repressor/activator), we first isolated and characterized a strong constitutive σ^A^ promoter from *B. megaterium*. Previously, a strong σ^A^ promoter was isolated from the pyruvate dehydrogenase (*pdh*) operon in *Bacillus subtillis* (39). For *B. megaterium*, we isolated the equivalent promoter from the *pdhABCD* (*bmd_1326-1329*) gene cluster by PCR along with the RBS of *pdhA* (*bmd_1326*) and used this to control the expression of *gfp* and mCherry. To initially validate the performance of gene expression *in vivo*, the optimised xylose-inducible promoter plasmid GFP variants (30) were used for comparison and tested under plate reader conditions. As expected, the growth rate of the empty vector control and non-induced xylose promoter recombinant strains was faster than the PDH or xylose-promoter strains, with all strains reaching the stationary phase after approximately 8 hours of growth (Supplementary Figure S1A). As a comparison between plasmid strains, normalised activity (RFU OD_600_^−1^ hr^−1^) for the PDH promoter was 1.4-fold higher than xylose-inducible promoter plasmid strains (Supplementary Figure S1B). For SDS-PAGE analysis, intracellular proteins were analysed from 100 mL batch-flask cultures for a qualitative assessment of GFP and mCherry production from xylose and PDH promoter plasmid strains (Figure 1A).

**Figure 1.**
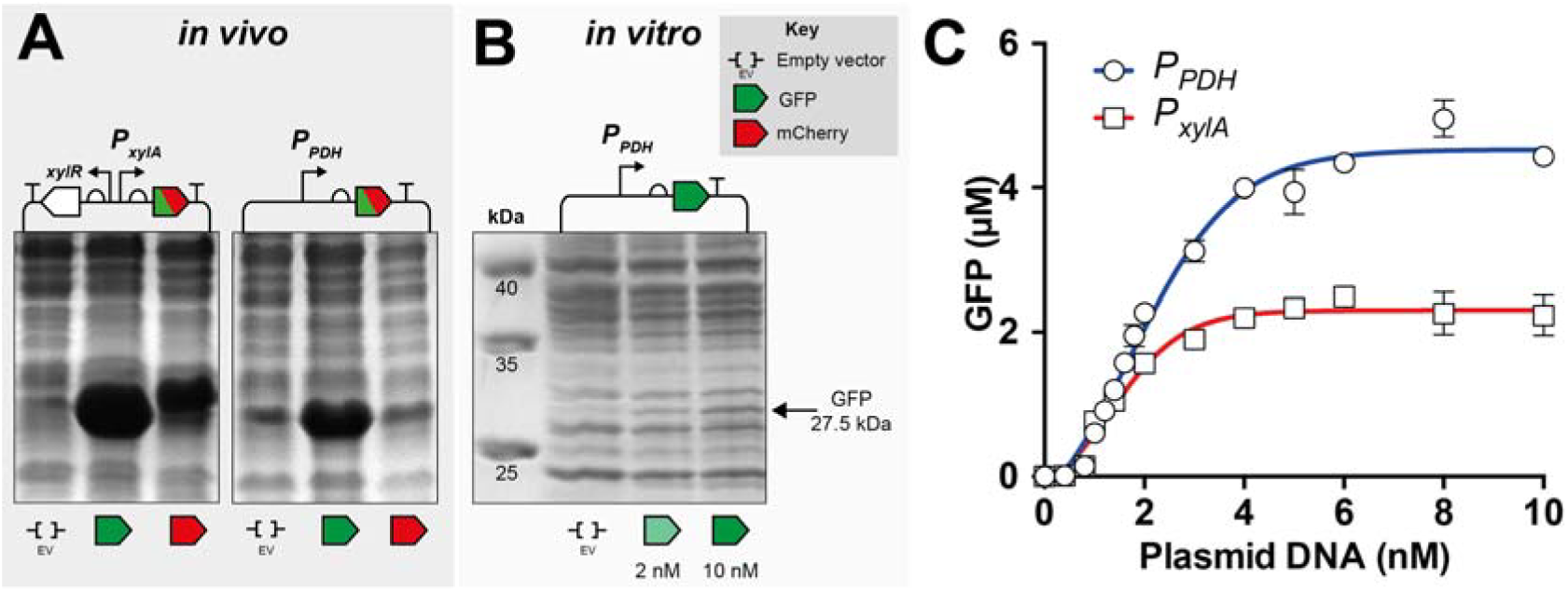
Comparison of the promoter systems in *B. megaterium* cells or ITT. **(A)** Recombinant *B. megaterium* DSM319 carrying the plasmids pSSBm85 (P_*xylA*_-*gfp*), pRBBm258 (P_PDH_-*gfp*), pSWBm19 (P_*xylA*_-*mCherry*) and pRBBm267 (P_PDH_-*mCherry*), respectively, were aerobically cultivated in LB medium. Intracellular proteins were analysed by SDS-PAGE. **(B)** SDS-PAGE analysis of *B. megaterium* cell-free incubations with either empty vector (EV – pMM1520) or pRBBm258 plasmid DNA. **(C)** End-point GFP values (μM) with increasing plasmid DNA concentration (nM) for the pRBBm258 and pKMMBm5 (lacking XylR repressor) plasmid.

### *B. megaterium* ITT activity

To test the xylose and PDH promoter systems *in vitro*, a *B. megaterium* cell-free extract was developed by using an *E. coli* S30 protocol for guidance (40). Extracts were prepared through four stages of preparation – growth, cell-disruption, heated incubation (“run-off” period) and dialysis. Following the S30 method with minor modifications (see Supplementary text) the cells were lysed with sonication followed by a 1.5 hour “run-off” period at 37°C and 3 hour dialysis. From this, significant levels of GFP synthesis were detected with 10 mg mL^−1^ of protein extract incubated with 10 nM of plasmid DNA, 1.5 mM amino acids and a 3-PGA energy buffer system. From this initial test, 1.4 μM of GFP was produced within 3 hours at 30°C. To optimise the method further, the sonication, “run-off” and dialysis stages of extract preparation were individually tested. Maximum activity was observed with an energy input of 154 kJ mL^−1^ for sonication and a “run-off” period of 1 hour at 37°C. These two modifications increased GFP batch synthesis up to 2.8 μM (Supplementary Figure S2). In addition, we also tested alternative incubation temperatures and time periods, but found the 1 hour incubation at 37°C to be optimal. Further details of optimisation are outlined in the Supplementary text. As a third preparation step, dialysis is used to remove small molecules, however, it is not always essential for the preparation of all *E. coli* cell-free extracts (41, 42). For *B. megaterium* DSM319, dialysis did not significantly affect ITT activity and therefore we consider it is not essential for this specific extract preparation. With further optimisation of Mg-glutamate (2 mM) and K-glutamate buffer (50 mM) conditions with 3-PGA as the energy source, maximum levels of 4.96 μM GFP (0.14 mg mL^−1^) are produced from 10 nM pRBBm258 PDH constitutive plasmid, with a band corresponding to GFP observable on SDS-PAGE (Figure 1B). This represents a 3.5-fold improvement from the initial test condition. With the xylose-inducible plasmid lacking the repressor XylR (pKMMBm5), maximum GFP batch synthesis of 2.48 μM is observed (Figure 2C). Maximal levels of protein production were saturated at approximately 5 nM of plasmid DNA for both promoter systems (Figure 1C). For comparison, fully optimised *E. coli* ITT systems can produce up to 80 μM (2.3 mg mL^−1^) of GFP in batch synthesis (1, 43). We also tested mCherry production *in vitro*, but production was limited to 0.17 μM under optimised conditions (Supplementary Figure S3 and Figure S4). This could be due to a high rare codon frequency for the *mCherry* gene (see Supplementary text).

**Figure 2.**
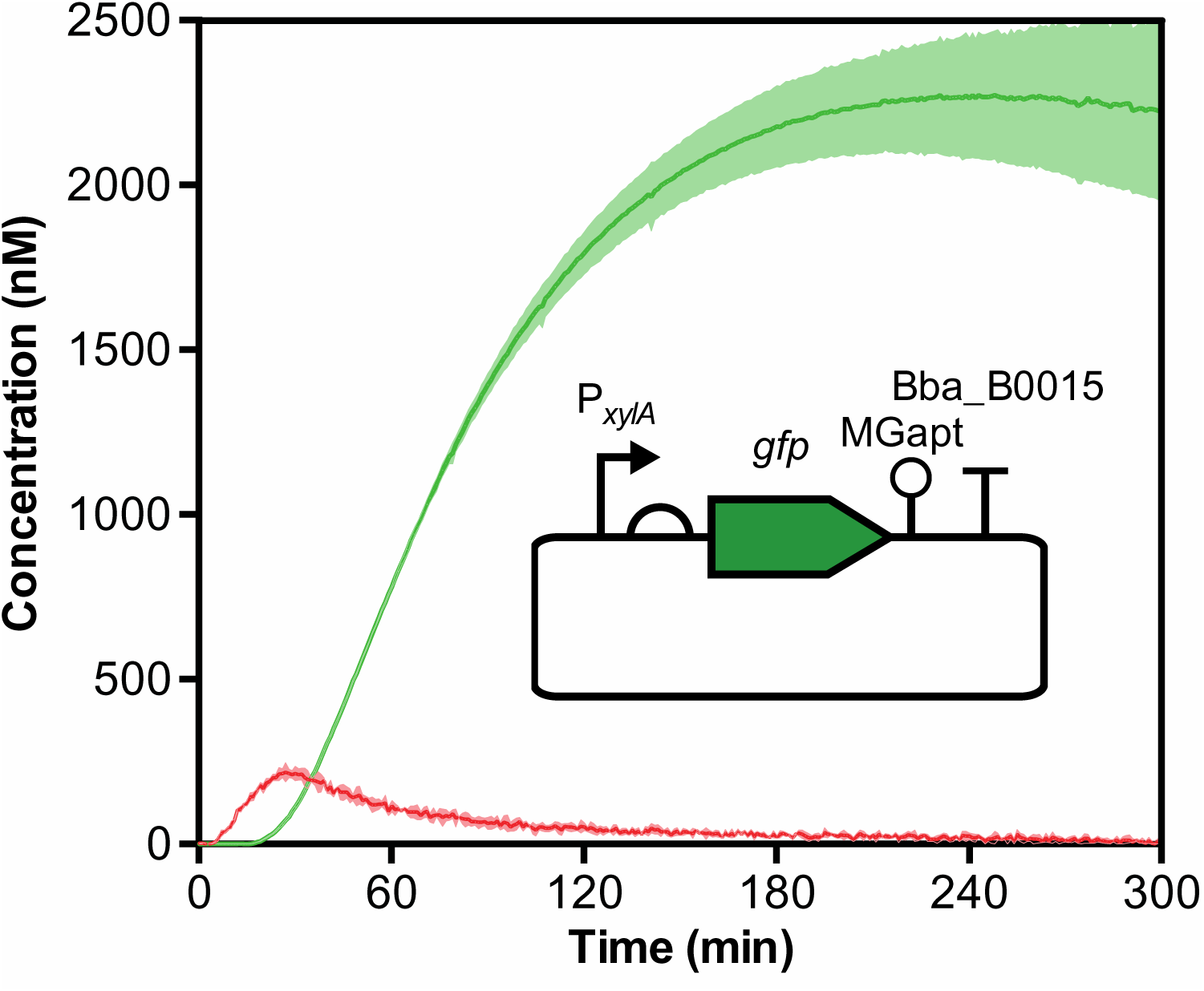
*B. megaterium* real-time ITT time course of mRNA and protein production. Design and experimental fluorescence time-course of 10 nM pKMMBm5-MGapt plasmid with mRNA (red line) and GFP (green line) reporters under standard conditions with 3-PGA energy buffer at 30°C. Measurements recorded every minute. Standard deviation is representative of a triplicate technical repeat and is displayed as a shaded area for each curve.

### Real-time mRNA and protein monitoring of the xylose-repressor system *in vitro*

The xylose-inducible promoter is derived from the native *B. megaterium xylABT* operon, which encodes the xylose isomerase (*xylA*), xylulokinase (*xylB*) and permease (*xylT*) genes (31, 44). Expression of the *xyl* operon is repressed from the divergently transcribed repressor XylR. In the absence of xylose, XylR can bind to two operator sequences (O_L_/O_R_) that overlap by 4 bp providing tight transcriptional gene expression control. Xylose binding to XylR releases the protein to allow transcription to start with gene expression induced up to 350-fold in *B. megaterium* DSM319, thus acting as a sensing mechanism for the cell to switch metabolism and utilise xylose as a carbon source (31). To study this system in cell-free reactions, we utilised a xylose reporter plasmid that lacks the XylR regulator (pKMMBm5). We decided to utilise the standard 3-PGA buffer for energy regeneration. Previously, this plasmid was shown to constitutively produce GFP at similar levels in wild-type *B. megaterium* DSM319 and WH325 Δ*xylR* mutant strains (45). This demonstrated that under standard growth conditions, the levels of XylR produced from the genomic copy of the gene were insufficient to repress *P*_xyl_ controlled GFP expression from the medium copy plasmid pKMMBm5 (45).

In order to characterize the kinetics of the xylose promoter in the absence of the repressor XylR, we wanted to use our cell-free system to simultaneously monitor transcription and translation *in vitro* (Figure 2). To monitor mRNA expression, we modified the 3’ untranslated region (UTR) to include a malachite green aptamer (MGapt) for real-time fluorescence detection of mRNA synthesis as previously described (46–51) (Supplementary Figure S5). mRNA degradation was estimated by measuring fluorescence for 2 hours and the order of mRNA degradation followed a single-phase exponential decay rate (Supplementary Figure S6 and Figure S7). A half-life estimated at 15.6 min is similar (18 min) to previously observed *E. coli* cell-free measurements (7).

We next tested a range of DNA concentrations in cell-free reactions with simultaneous measurement of MGapt and GFP fluorescence. With increasing DNA concentration, the rate of MGapt fluorescence rises rapidly within 6 min up to a peak concentration of approximately 235 nM of GFP-MGapt transcript at 25-27 min after the start of incubation. Thereafter, the signal decays suggesting that either nucleotide or energy regeneration becomes rate limiting for further mRNA synthesis. Spiking the reaction with a purified GFP-MGapt transcript compared to the equivalent plasmid control, shows that mRNA levels are more stable throughout the assay when expressed from plasmid DNA, suggesting that continuous transcription provides mRNA substrate for translation throughout the reaction time-period (Supplementary Figure S8).

### Bayesian modelling of the xylose-repressor system

To characterize the behaviour of the xylose-repressor system in more detail, we chose to model the system using ordinary differential equations (ODEs) with 4 species and 15 parameters (Supplementary Tables S1 and S2). These parameters were simultaneously inferred from multiple experimental data sets using a Bayesian Markov chain Monte Carlo (MCMC) approach (see Methods for details). A posterior distribution of the model parameters was inferred from the GFP and mRNA time course data sets with the DNA template (pKMMBm5-MGapt) titrated at a range of different concentrations. These parameters were simultaneously inferred from multiple experimental datasets and chemical species (mature GFP and mRNA) using a combined log-likelihood function (Supplementary equation S1). Multiple MCMC runs were generated from different parameter starting points and convergence was confirmed for all parameters using the Gelman-Rubin diagnostic(52). Simulated trajectories from the model closely matched the experimental time-course data for both GFP and mRNA concentration at different DNA template concentrations (Figure 3 and Supplementary Figure S9 and Figure S10), whilst the posterior distribution was found to be unimodal (Supplementary Figure S11). The reaction machinery appeared to be saturated at around 5 nM template DNA with no further increase in total protein produced, however, mRNA expression continued increasing with higher DNA concentrations. This implied that the ribosomes were saturated with mRNAs but the host RNA polymerase was not saturated with DNA template.

**Figure 3.**
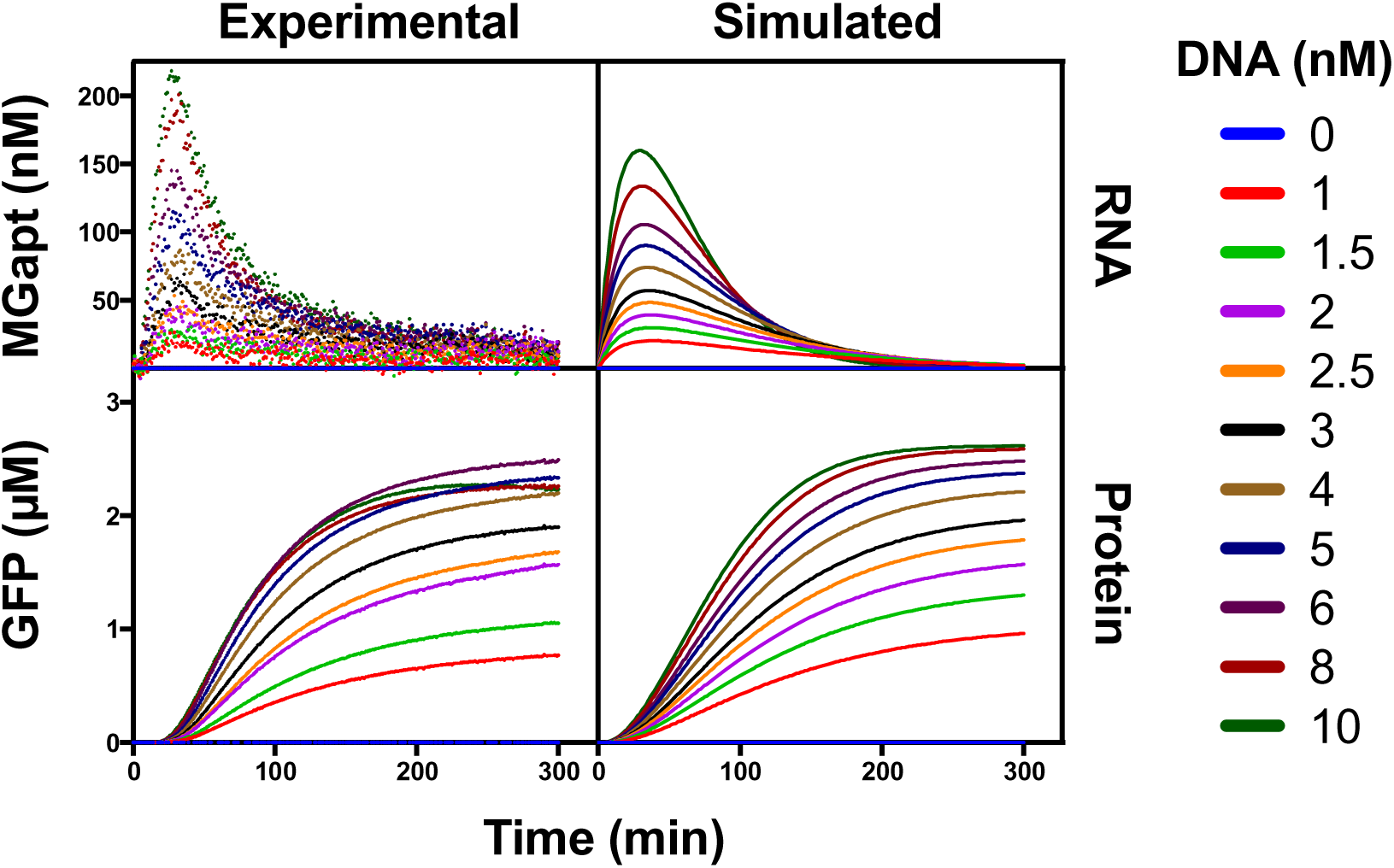
Transcription and translation of the xylose-inducible promoter. Cellfree extracts (10 mg mL^−1^) were incubated at 30°C for 6 hours with a range of concentrations (1-10 nM) of the pKMMBm5-MGapt plasmid. Fluorescence data was collected every 60 seconds for MGapt (mRNA) and GFP (protein) signals. Experimental data were modelled using ordinary differential equations with a system of 4 species and 15 parameters detailed in equation 1 and Table S1 and 2, and parameter were inferred using Markov chain Monte Carlo.

### Liquid handling robotics of cell-free reactions for augmented full factorial inducible promoter characterization

A second, larger-scale experiment was carried out on the XylR repressible promoter to separately characterize its kinetics in cell-free reactions. This experiment was designed to simultaneously monitor GFP translation with 108 unique conditions in triplicate (324 total reactions) made up of an augmented full factorial combination of different concentrations of purified recombinant XylR repressor protein (0-1000 nM; Supplementary Figure S15), D-xylose (0-1000 μM), and pKMMBm5-MGapt DNA template (1, 2 and 5 nM) on a 384 well microtiter plate using an acoustic liquid handling robot (Figure 4A).

**Figure 4.**
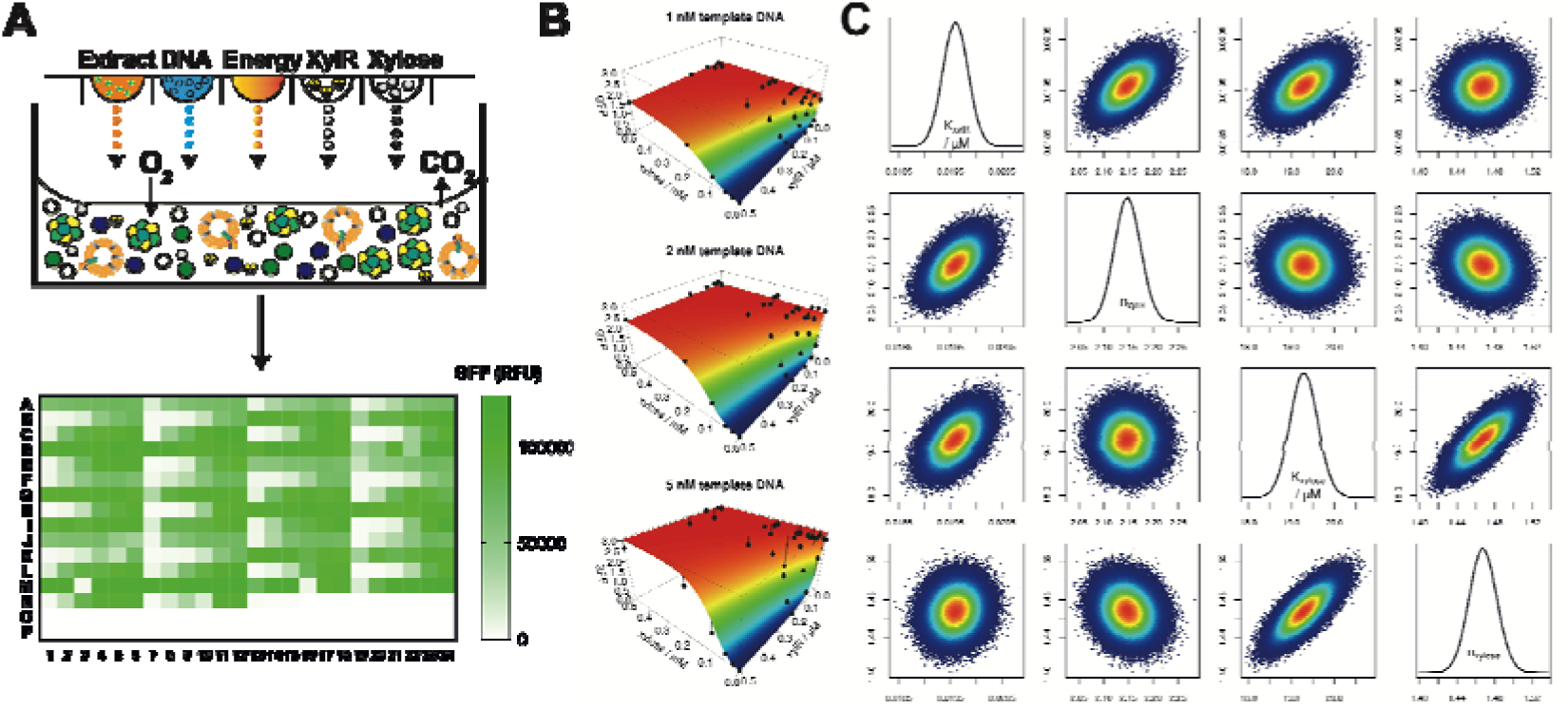
Cell-free prototype validation and experimental augmentation of the xylose-inducible promoter system. Experimental and Bayesian modelling characterization of the xylose-inducible promoter system in cell-free. **(A)** Dxylose (mM), purified XylR^His^ (μM) and pKMMBm5 plasmid DNA (nM) was titrated in a cell-free reaction using an Echo^®^ 525 acoustic liquid handling system (LabCyte) with 25 nL transfer droplets. The end-point value of GFP is provided for this experimental dataset as a surface heat map for 324 conditions. Experimental data was simultaneously modelled with 4 species and 15 parameter variables with MCMC analysis. **(B)** The maximum likelihood simulated end-point GFP concentrations are shown as a surface with the experimental end-point GFP concentrations shown as black points. The distance between the modelled surface and the experimental values is shown as a connecting black line. **(C)** The inferred posterior distribution over the K_D_ and Hill-coefficients for binding of XylR and xylose.

In the absence of D-xylose and XylR^His^, GFP is constitutively produced and slightly elevated by the addition of 0.5 mM D-xylose. We also verified that the *B. megaterium* ITT system is not inhibited by high D-xylose concentrations (Supplementary Figure S16). When XylR^His^ is titrated in the absence of D-xylose, a repression effect on GFP production is observed, as expected, but this effect is variable with DNA concentration (Figure 4B). Following the titration of D-xylose in the presence of 0.1 to 1.0 μM XylR^His^, GFP expression is recovered, which is consistent with the release of the XylR^His^ repressor from the promoter region upon binding of D-xylose (Figure 4B).

Kinetic data from the first set of experiments was incorporated into the model by fitting a multivariate Gaussian distribution to the posterior distribution of the first experiment (Supplementary Figure S11). This was then used as the prior distribution for inferring model parameters using data from the second set of experiments with the same MCMC method (Fig 4C, Supplementary Tables S1-2 and Figure S17). Over this diverse range of conditions, both the simulated final GFP concentrations (Figure 4B) and the time-course trajectories were found to fit the experimental data remarkably well (Supplementary Figure S12-14). The maximal translation rate was inferred to be approximately 27-47 nM (GFP) min^−1^, which is similar to previously determined figures for *E. coli* cell-free reactions (53), whilst the maximal transcription rate was inferred at between 310-380 nM (mRNA) min^−1^. Previously, the maturation rate of GFP+ (a synonym for GFPuv3) was measured to be 0.066 min^−1^ (54). We independently infer the maturation rate to between 0.047-0.057 min^−1^. The difference may reflect inaccuracies in the model or a lower oxygen concentration could limit maturation in the reaction mixture. Moreover, the XylR repressor was found to cooperatively bind to its operator sequence (Hill coefficient 2.1-2.2), while xylose appears to cooperatively bind XylR (Hill coefficient 1.4-1.5).

### Succinate as an alternative substrate for energy regeneration

For future scaled-up cell-free protein production in *B. megaterium*, a minimal, low-cost energy system for extract preparation is required (55). For *B. megaterium* cell-free we initially investigated the importance of each individual component (42) in the standard energy buffer for *B. megaterium* ITT activity. Firstly, in the absence of NTPs, neither mRNA nor GFP is synthesised. Interestingly, if tRNA’s, NAD^+^, coenzyme A and folinic acid were removed, this did not significantly affect ITT activity in *B. megaterium* extracts (Supplementary Figure S18). In addition, the removal of 3-PGA only led to a 75% reduction in GFP synthesis, whilst spermidine and cAMP were considered non-essential, but beneficial for activity. Next, we screened potential energy substrates including glucose, maltose, PEP, pyruvate, succinate and glutamate to replace 3-PGA as a sole energy source. The residual activity from the absence of 3-PGA is likely to be due to K-glutamate or other possible amino acid catabolic pathways. Glutamate has previously been shown as a low-cost energy source in *E. coli*, where it is enters the Krebs cycle through α-ketoglutarate (42). For alternative energy sources in *B. megaterium*, we observed strong activity for pyruvate and succinate, however, no activity was observed with glucose, maltose or phosphoenolpyruvate (Supplementary Figure S19). Focusing on succinate, we observed a strong response in GFP production in response to an increasing concentration of the substrate, which peaked in activity at 50 mM Na-succinate (Supplementary Figure S20). This effect was also pH-dependent (Supplementary Figure S21) with maximal activity of GFP synthesis observed at pH 8.2, although RNA synthesis appeared to optimal at lower ranges around pH 7.0.

## Discussion

There is a growing interest in exploring arcane microbial platforms for future cellular factories (13). However, the advantages of using such hosts are often counter balanced by a lack of genetic tools, poor genetic tractability or fastidious growth. Generally to engineer bespoke hosts for a specific industrial process, classical strain evolvement by mutagenesis is utilised to optimise production of fine chemicals, amino acids and vitamins (23).

Recently, cell-free ITT platforms have been established for prototyping genetic parts and testing metabolic pathway designs (1, 6, 49). Here we have utilised the advantages of cell-free ITT to establish a *B. megaterium* cell-free platform for prototyping genetic parts and pathway designs for a host that is difficult to engineer but has the potential for industrial scale catalysis of low-cost building blocks (e.g. crude glycerol, sugar beet) into value-added products (20, 56). Moreover, there is a general opportunity in synthetic biology to utilise alternative chassis for bespoke applications, for example accessing challenging recombinant enzymes (25, 36).

To provide a robust assessment of our *B. megaterium* cell-free platform, we also describe a factorial experimental design and modelling approach to delineate the kinetic behaviour cell free reactions through automated experimental testing and Bayesian statistical parameter inference. To our knowledge, our use of automation and Bayesian modelling are the first time this has been applied to cell-free prototyping thereby generating complex datasets for refined and accurate analysis allowing the design of robust synthetic genetic circuits using Bayesian design methods (57). In order to test our cell free extract preparations we isolated and characterized a *B. megaterium* specific σ^A^ constitutive promoter along with the xylose-inducible promoter. In comparison to a fully optimised *E. coli* cell-free reaction using σ^70^ constitutive promoter or T7 promoter (43), our initial *B. megaterium* extracts produce approximately 10-fold less GFP using the σ^A^ promoter. Significantly, the GFP production capacity of *B. megaterium in vivo* is comparable (gram per litre scale) to optimised *E. coli* strains (58, 59). This suggests that the current productivity of our cell-free system is below the true potential for this host. From our experimental and modelling data, mRNA expression is not rate limiting but the translation apparatus appears to be saturated above 5 nM of plasmid DNA. To optimise gene expression in cell-free systems, current strategies have focused optimising energy regeneration (60), mRNA expression (61) or ribosome enrichment (62, 63). Therefore for the moment at least, it would seem that either the aminoacyl-tRNA synthetases or ribosome activity is rate limiting for *B. megaterium* ITT and therefore a key starting point for further investigation. Another unknown factor is the availability or stability of the tRNA derived from the host extract. However for the purpose of characterizing our *B. megaterium* ITT system the expression levels of our test plasmids give strong and discernable signals.

Efficient energy regeneration is an essential component of a cell-free extract to replenish ATP and nucleotide supplies for protein synthesis from DNA (60). To our knowledge we have established for the first time that succinate can act as a substrate for energy regeneration in cell free extracts (Figure 5). Succinate dehydrogenase in bacteria is found as a three-component (ShdABC) membrane bound complex and is unique since it is the only Krebs cycle enzyme step that also participates in the electron transport chain as part of respiratory complex II. The *Bacillus subtillis* SdhABC complex has previously been characterized by electron paramagnetic resonance and shown to catalyse a one-step oxidation of succinate to fumarate, using an electron tunnelling mechanism (64). It has been proposed that inverted vesicles derived from the inner membrane of *E. coli* form during cell-free extract preparation and contain several active members of the electron transport chain and ATP synthase to regenerate energy through oxidative phosphorylation (65). Our observed ITT activity with 3-PGA, glutamate and pyruvate suggest that inverted vesicles are also likely to form in *B. megaterium* cell-free extracts. The use of succinate provides a potential low-cost route for scaling-up *B. megaterium* cell-free systems in the future, which is an essential requirement of cost-efficient production methods (55).

**Figure 5.**
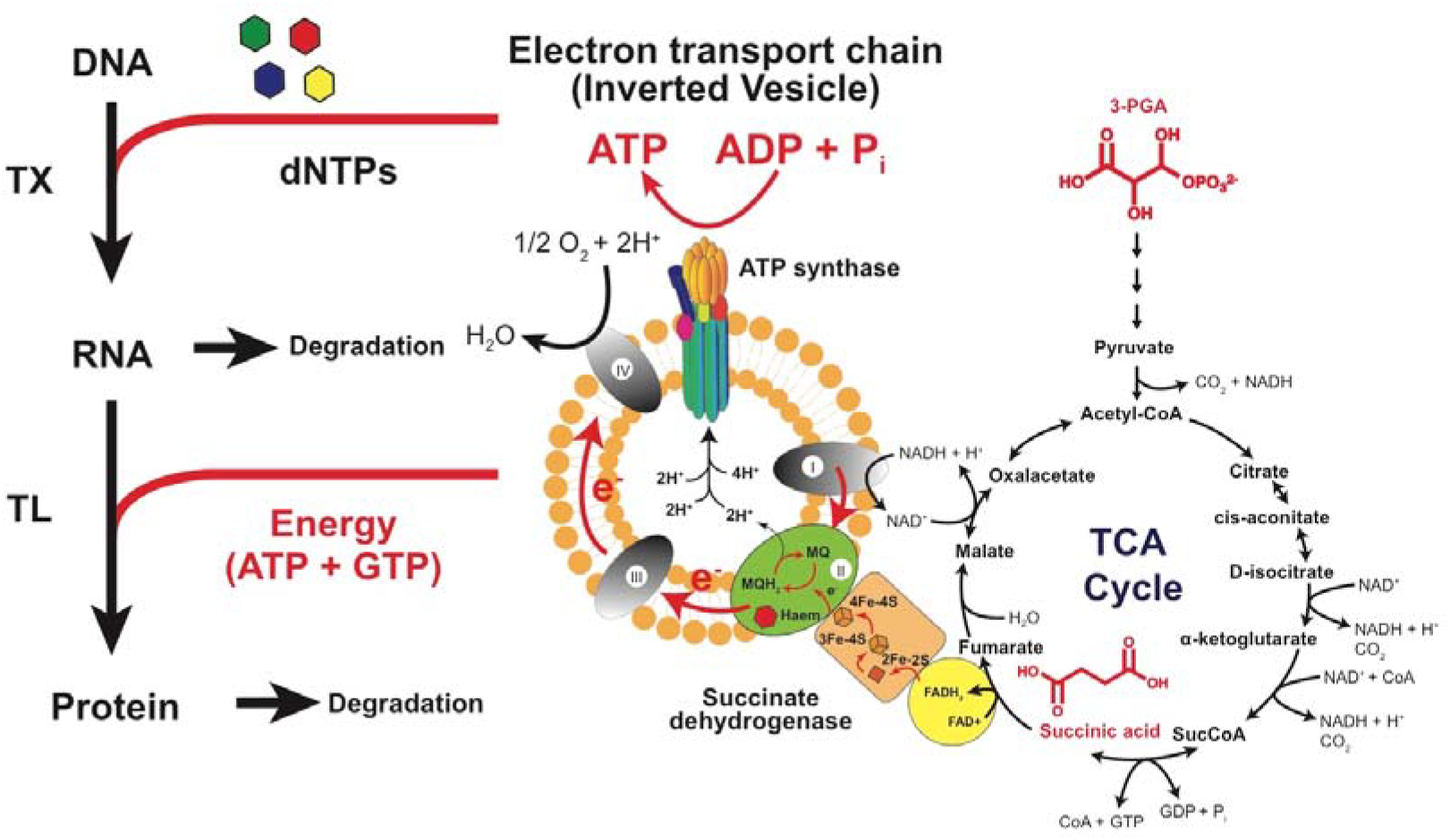
Energy regeneration and ITT reaction components in *B. megaterium*. A route for energy regeneration through either 3-PGA or succinic acid as energy substrates for ATP synthesis. Theoretical model of an inverted vesicle harbouring the electron transport chain for providing energy for transcription and translation stages.

To begin studying genetic circuits in *B. megaterium* cell-free we investigated the xylose-repressor system, which switches on gene expression in response to xylose. Aside from providing high-inducible levels of expression in the presence of D-xylose, this XylR based regulatory system provides strong trans regulatory control in the absence of xylose (31). Interestingly, by reconstituting this system *in vitro* using purified XylR, we also observe tight regulatory control of GFP expression, with our Bayesian model estimating the binding K_D_ for XylR and xylose at 19.6 nM and 19.3 μM, respectively. This demonstrates a simple reconstitution and modelling characterization of an inducible regulatory element in *B. megaterium* cell-free extracts and thus provides a basis for future prototyping of *B. megaterium* specific regulatory elements. These rigorously characterized elements can then be rationally composed into more complex circuits (7, 47, 57).

Cell-free synthetic biology is a growing area of research for rapid part and pathways design prototyping to large industrial scale protein manufacturing (55). Our demonstration of *B. megaterium* as new host for cell-free synthesis using automated nanolitre scale liquid robotics combined with a rigorous factorial experimental design and Bayesian modelling approach opens up the possibility of systematically exploring less-well studied chassis for cell-free synthetic biology applications.

## Acknowledgements

SW and RB would like to thank Dieter Jahn for support and discussion.

